# A novel mouse model for developmental and epileptic encephalopathy by Purkinje cell-specific deletion of *Scn1b*

**DOI:** 10.1101/2024.11.19.624370

**Authors:** F. Isaac Guillén, Mendee A. Geist, Shao-Ying Cheng, Arwen M. Harris, Martha E. Treviño, Audrey C. Brumback, Hiroshi Nishiyama, MacKenzie A. Howard

## Abstract

Loss of function variants of *SCN1B* are associated with a range of developmental and epileptic encephalopathies (DEEs), including Dravet syndrome. These DEEs feature a wide range of severe neurological disabilities, including changes to social, motor, mood, sleep, and cognitive function which are notoriously difficult to treat, and high rates of early mortality. While the symptomology of *SCN1B*-associated DEEs indicates broad changes in neural function, most research has focused on epilepsy-related brain structures and function. Mechanistic studies of *SCN1B*/*Scn1b* have delineated diverse roles in development and adult maintenance of neural function, via cell adhesion, ion channel regulation, and other intra- and extra-cellular actions. However, use of mouse models is limited as knockout of *Scn1b*, globally and even in some cell-specific models (e.g., Parvalbumin+ interneuron-specific knockout) in adult mice, leads to severe and progressive epilepsy, health deterioration, and 100% mortality within weeks. Here, we report findings of a novel transgenic mouse line in which *Scn1b* was specifically deleted in cerebellar Purkinje cells. Unlike most existing models, these mice did not show failure to thrive or early mortality. However, we quantified marked decrements to Purkinje cell physiology as well as motor, social, and cognitive dysfunction. Our data indicate that cerebellar Purkinje cells are an important node for dysfunction and neural disabilities in *SCN1B*-related DEEs, and combined with previous work identify this as a potentially vital site for understanding mechanisms of DEEs and developing therapies that can treat these disorders holistically.

## Introduction

*SCN1B* (sodium voltage-gated channel beta subunit 1) variants have been reported in people with a range of neurological and cardiac disorders. Full autosomal loss of function causes Developmental and Epileptic Encephalopathies (DEEs), including Dravet syndrome (Wallace et al., 1998; Ogiwara et al., 2012; Aeby et al., 2019). DEEs are catastrophic pediatric disorders characterized by neurodevelopmental disabilities that can include disorders of motor, social, mood, sleep, cognitive function, and medically refractory epilepsy. While these conditions are characterized by widespread neurological dysfunction, most research on *SCN1B* and other DEEs focuses on epilepsy and seizure-related phenotypes and brain regions. Importantly, the non-seizure-related disabilities often persist or continue to worsen even when seizures are medically controlled or as seizure burden decreases with age. These non-seizure neurological disabilities are reported by many caregivers as having the worst impact on quality of life (Villas et al., 2017).

*SCN1B* encodes two splice variant proteins, β1 and β1B, that each play multiple roles in development and physiology. Each can act as an auxiliary ion channel subunit that traffics and modulates the physiology of voltage-gated Na^+^ (Isom et al., 1992), K^+^, and likely other ion channels (Marionneau et al., 2012). Each also acts as a cell adhesion molecule and plays a role in axon guidance and developmental patterning (Brackenbury et al., 2008, 2010, 2013; Lopez-Santiago et al., 2011; Patino et al., 2011). Recent work shows that β1 can be cleaved and a fragment can enter the nucleus where it regulates gene transcription (Bouza et al., 2021). In keeping with its range of functions, loss of *Scn1b* causes a mix of cell type-specific changes: altered patterning and axonal fasciculation (Brackenbury et al., 2013), hyperexcitability in some but not all types of forebrain pyramidal neurons (Marionneau et al., 2012; Reid et al., 2014; Hull et al., 2020; Chancey et al., 2023), hypoexcitability in cerebellar granule cells (Brackenbury et al., 2010), hypoexcitability in some but not all subtypes of GABAergic interneurons, and changes to synaptic integration (Hull et al., 2020; Chancey et al., 2023). While such phenotypes will intuitively cause major disruptions to overall neural function, linking specific cellular level changes with neurological disabilities of *SCN1B*-DEE is difficult: *Scn1b*-/- mice have severe and progressive health deterioration and seizures starting in the second postnatal week, leading to 100% mortality by postnatal day 22 (Chen et al., 2004). Further, excision of *Scn1b* from forebrain neurons in adult mice results in 100% mortality within three weeks (O’Malley et al., 2019). This illustrates the vital role of *Scn1b* in the maintenance of neural function beyond development, but also impedes the study of neurological deficits that affect patients.

Our goal was to develop and validate a *Scn1b* loss of function mouse model robust enough to allow study of mechanisms for a range of DEE-associated neurological disabilities. An understudied point of convergence for altered neurological function in DEEs is the cerebellum, which is known to be involved in motor control, cognition/memory, and social behavior (Reeber et al., 2013). Importantly, cerebellar granule cells and Purkinje cells (PCs) show high expression of *SCN1B* (Lein et al., 2007) and *Scn1b* has been shown to play a role in the specialized physiology of these neurons (Aman et al., 2009). Thus, toward our goal, we crossed a floxed *Scn1b* knockout mouse (Chen et al., 2007) with a Purkinje cell-specific Cre line (Barski et al., 2000). These Purkinje cell-specific *Scn1b* knockout mice survived and thrived into adulthood, but showed profound deficits in PC excitability, locomotion, social behavior, and cerebellar associative learning. The data presented here provide insights into potential mechanisms for *SCN1B* function in the developing and mature brain and DEE-related disabilities.

## Material and Methods

### Animals

All experiments were performed in accordance with protocols approved by the Institutional Animal Care and Use Committee of University of Texas at Austin. All mice were housed in standard group housing on a 12:12 light cycle, provided food and water *ad libitum*. All mice were strictly maintained on a congenic C57Bl/6J background, with regular back-crossings with C57Bl/6J (strain #000664) mice purchased from Jackson Labs to eliminate genetic drift. We performed tail/toe clipping at postnatal day (P)6-8 for identification. Genotyping was performed on tail tissue samples by Transnetyx using real-time PCR.

Global-*Scn1b* knockout (KO) mice, i.e., *Scn1b*-/-, were a generous gift from Lori Isom at the University of Michigan (Chen et al., 2004). *Scn1b*+/- breeding pairs produced experimental mice (Global-*Scn1b* KO) and wild-type littermates, used for experiments as juveniles at P15-20. Global-*Scn1b* KO mice in our colony experienced spontaneous seizures and stunted growth beginning around P10 and 100% mortality by P22 as previously reported (Chen et al., 2004; Hull et al., 2020, Chancey et al., 2023).

PC-specific Cre mice (L7/*Pcp2*-Cre) were obtained from Jackson Labs (strain #004146). PC-specific *Scn1b* KO mice used in this study were generated from breeding pairs of female *Scn1b^flox/flox^*;Ai14-TdTom reporter positive (Jackson Labs, Strain #007914);L7/*Pcp2*-Cre positive mice, crossed with male *Scn1b^flox/flox^*, Ai14-TdTomato reporter positive mice. Experimental mice (hereafter referred to as PC-*Scn1b* KO) were male and female *Scn1b^flox/flox^*;Ai14-TdTom+;Cre+. Control mice for these experiments (hereafter referred to as PC-WT) were *Scn1b^flox/flox^*;Ai14-TdTom+;Cre- male and female littermates. Experiments with these mice were performed in juveniles at P15-20 (2.5 wk of age), or in adults P56-88 (8-12 wk of age).

### Preparation of Acute Cerebellar Slices

Mice were anesthetized with a mix of ketamine (90 mg/kg) and xylazine (10 mg/kg) intraperitoneally followed by a transcardial perfusion with ice-cold oxygenated cutting solution, containing (in mM): 205 sucrose, 25 sodium bicarbonate, 2.5 KCl 1.25 sodium phosphate, 7 MgCl_2_, 7 D-glucose, 3 sodium pyruvate, 1.3 ascorbic acid, and 0.5 CaCl_2_. Brains were rapidly removed, and parasagittal slices of cerebellar vermis/paravermis (250 µm) were cut with a vibratome (Leica VT 1200) filled with oxygenated cutting solution. Slices were incubated for 30 min at 37°C in holding solution made of (in mM): 125 NaCl, 25 sodium bicarbonate, 2.5 KCL, 1.25 sodium phosphate, 12.5 D-glucose, 2 MgCl_2_, 2 CaCl_2_, 1.3 ascorbic acid, and 3 sodium pyruvate, and then at room temperature for at least 30 min before transfer to the recording chamber.

### Whole-Cell Current Clamp Recordings

Recordings were obtained in juvenile (2.5 wk) and adult (12 wk) mice cerebellar PC somata in acute slices using a Multiclamp 700B amplifier interfaced with a Digidata 1550B acquisition system and pClamp 11.1 software (Molecular Devices). Slices were continuously bathed in warm (32.5°C) oxygenated artificial cerebrospinal fluid (ACSF), containing (in mM): 125 NaCl, 25 NaHCO_3_, 2.5 KCl, 1.25 NaH_2_PO_4_, 12.5 D-glucose, 1 MgCl_2_, and 2 CaCl_2_. Patch pipettes of 2.5 - 3.5 MΩ were pulled from borosilicate filamented glass (Sutter BF150-110-10) using a P-1000 puller (Sutter). Pipettes were filled with a potassium gluconate solution containing (in mM): 120 K-gluconate, 20 KCl, 10 HEPES, 4 NaCl, 1 EGTA; 4 Mg-ATP, 0.3 Na-GTP, and 7 phosphocreatine disodium salt hydrate. Pipette capacitance neutralization and bridge balance were applied. Spontaneous firing activity was recorded in the absence of injected current for 20 seconds. In the evoked spiking and voltage sag protocols we applied holding current to stop PC spontaneous firing and give the neurons a baseline membrane potential of ∼-65 mV. Evoked spiking was generated by depolarizing current injections from 0 pA to 600 pA with steps of 25 pA, 1 second duration. Voltage sag and input resistance were measured with hyperpolarizing current steps from 0 to -600 pA in 50 pA steps of 1 second duration. In the spike intrinsic properties protocol, we used the first spike out of the train of spikes evoked at 300 pA current injection of 1 second duration. Data were acquired at 50 kHz.

### Physiology analysis

We identified the action potential threshold (in mV) by “change in dV/dt”, using the first derivative of the voltage trace and took the standard deviation (SD) of the dV/dt milliseconds before the action potential (Atherton and Bevan, 2005; Howard and Rubel, 2010). The membrane potential threshold was reached once the dV/dt reached: mean_dV/dt_ +(6*SD_dV/dt_), relative to the dV/dt baseline measured 4 milliseconds before the action potential. We also visualized the action potential threshold as the inflection point of a phase plane plot (Fig. 3A, bottom images) using changes of the membrane potential with time (dV/dt measured as mV/ms; y-axis) plotted against the membrane potential (measured as mV; x-axis) (Jenerick, 1963). The action potential peak was taken as the maximum membrane potential after threshold, and the action potential amplitude as the difference between threshold and peak. The action potential width was measured at the half-amplitude of the action potential. The after-hyperpolarization was identified as the minimum membrane potential right after the spike peak voltage followed by the repolarization phase (Fig. 3A, top images). Data were not corrected for junction potential.

### Histology

Cerebellum size and PC number were analyzed in adult (12 wk) PC-*Scn1b* KO and PC-WT mice. Mice were deeply anesthetized and then perfused with phosphate buffered saline (PBS) followed by 4% paraformaldehyde in PBS. After 24 hr fixation, sagittal slices were prepared for immunohistochemistry as previously described (Howard et al., 2014). Vermis slices were incubated with an antibody to Calbindin (Calb; monoclonal anti-Calbindin-D-28K; Sigma catalog # C9848; 1:500), a highly selective marker for PCs (Garcia-Segura et al., 1984). Slices were then incubated with secondary goat anti-mouse IgG antibodies conjugated to fluorophores (Alexa 594; Thermofisher catalog # A- 21235; 1:500). Mounted slices were imaged on a Nikon AR-1 confocal microscope in the Center for Biomedical Research Support Microscopy and Imaging Facility at UT Austin (RRID:SCR_021756).

Cerebellum size and PC number were quantified using only the Calb channel (green) so that researchers remained blinded to genotype. To measure size, the cerebellum was outlined, and area was calculated using ImageJ (RRID:SCR_003070). For consistency, the same region of cerebellum (the ventral aspect of the 9^th^ lobule of the cerebellar vermis) was used to quantify PC density in each brain. A line was drawn in ImageJ along the PC cell body layer, measuring at least 1.2 mm, and the number of identifiable PCs lying along this line was counted. Once this blinded analysis was completed, the same line was used to count PCs in the Cre positive channel (magenta) of the image to quantify the number of labeled PCs that also expressed Cre-driven TdTomato.

### Behavior

Mice used for behavior experiments were kept on a reverse light:dark cycle, all behavior experiments were performed during the dark cycle. Mice were allowed to acclimate to the behavior room for 30 minutes prior to handling habituation and behavior testing. Handling habituation was carried out in the behavior room for 5 days, 5 minutes each day. On habituation days 4 and 5, mice were also habituated for 5 minutes in the three-chamber arena. For behavior testing, open field and three-chamber tasks were performed on consecutive days, and repeated when mice were P56-60 (8 wk), P70-74 (10 wk), and P84-88 (12 wk). A digital camera and ManyCam version 8.1.0.3 video software (Paltak, Inc. Jericho, NY) was employed for mouse tracking. The behavior room was illuminated with red A19 LED light bulbs and the computer monitor was covered with a red cellophane sheet.

Open field behavior was measured in a neutral white Plexiglass box (L50 x W50 x H25 cm). The mouse was placed in the center of the square arena and allowed to explore freely while being recorded for 5 minutes. Total distance traveled and time spent in the center zone of the arena (20 x 20 cm) were calculated using EthoVision XT 16 (RRID:SCR_000441) software.

The three-chamber test was performed using a rectangular white Plexiglass box (L60 x W40 x H25 cm), divided into three chambers each 20 x 40 x 25 cm in size. The dividers had rectangular openings (10.5 x 25 cm) allowing mice free access into each chamber. During habituation to the chamber the dividers were removed. The three-chamber test was performed in three phases. In phase one, clean wadded paper towels were placed under wire cages in each side chamber. The test mouse was placed in the center chamber and allowed to freely explore all three chambers for five minutes. In phase two, the paper towels were replaced with a novel mouse under the wire cage in one side chamber, and by a novel object under the wire cage in the other side chamber. The novel mouse was sex- and age-matched to the test mouse. The test mouse was returned to the center chamber and again allowed to explore freely for 5 minutes. In phase three, the novel object was replaced with a new novel mouse to compare interest between the familiar mouse and the novel mouse; the test mouse was then allowed another 5-minute free exploration. The test mice were judged to be in a chamber when the center point of the mouse crossed through the divider. Time spent in each chamber, crossings into each chamber, and total distance traveled were quantified.

### Eyeblink Conditioning

Adult PC-*Scn1b* KO and PC-WT mice were used in delay eyeblink conditioning experiments. Mice were anesthetized with isoflurane (5% for induction and 1-2% for maintenance). Carprofen (10 mg/kg) was subcutaneously administrated as an analgesic. Lidocaine (7 mg/kg) was also subcutaneously administrated under the scalp as a local anesthetic. Then, the scalp was cut to expose the skull surface, and soft connective tissues on the skull were removed. A thin layer of surgical cyanoacrylate (Vetbond, 3M) was applied on the cleaned skull surface, and a custom-made thin titanium head plate was attached on top of the cyanoacrylate with dental cement (C & B Metabond, Parkell). After recovering from anesthesia, the mice were returned to the home cage. Carprofen (10 mg/kg) was administrated every 24 hours for 2 days post-surgery as an analgesic.

The overall setting and procedure for eyeblink conditioning were similar to a previous publication (Siegel et al., 2015). At least a week following the surgery, mice were allowed to acclimate to a custom-built rotating wheel. Each mouse was securely positioned on the wheel using the head plate and allowed to run freely for approximately 5 minutes (Day 1), 10 minutes (Day 2), 20 minutes (Day 3), and 30 minutes (Day 4).

Delay eyeblink conditioning began after 4 days of acclimation. Each mouse received 10 training sessions, with one session per day. A training session consisted of 18 blocks of 5 trials. One block consisted of four conditioning trials where we applied a conditioned stimulus (CS; blue LED) and unconditional stimulus (US; periocular air puff, 5 psi) and one probe trial (only CS). The CS duration was 280 milliseconds. It does not elicit eyelid responses in naïve animals. The US duration was 30 milliseconds and was applied 250 milliseconds after the CS onset; this way, the CS and US co-terminated. The US always elicits reflexive eyelid closure. An inter-trial interval was randomized between 5-15 seconds, with an average of 10 seconds. In each trial, eyelid closure was video recorded at 250 frames per second using an infrared light and a high-speed camera (Prosilica GE, Allied Vision). The training session was controlled by a custom-written IGOR Pro (RRID:SCR_000325) code.

To quantify the behavior, we measured the fraction of eyelid closure (FEC) and the conditioned response (CR) rate. FEC indicates how much an eyelid closed during a trial compared to the beginning of the trial. It ranged between 0 (eyelid opened) and 1 (eyelid closed) in most trials but occasionally became negative. The negative FEC indicates that the eyelid, which was partially closed at the beginning, widened before the US was delivered. Trials during which FEC dropped below -0.1 were discarded from analysis, but such trials consisted of only 3.0% of all trials. An eyelid closure was considered a CR if FEC was below 0.1 during the first 100 milliseconds of the CS presentation and above 0.1 immediately before the US presentation. These criteria are to distinguish CR from randomly occurring ill-timed eyelid closures. CR rate is defined as the ratio of CR-containing trials to all trials in a single training session.

### Data analysis and statistics

Electrophysiological data were analyzed using MATLAB 2022a (RRID:SCR_001622) software. We performed statistical analyses appropriate to the specific experiment and data set with GraphPad Prism 9 (RRID:SCR_002798). All figures show mean ± SEM and N = (cells/mice). For clarity, statistical tests and p values are reported within the text, while descriptive statistics and further details of statistical analyses are presented in **Table 1** for t-tests and **Table 2** for ANOVAs. Statistical significance is defined as p<0.05.

**Table 1:**
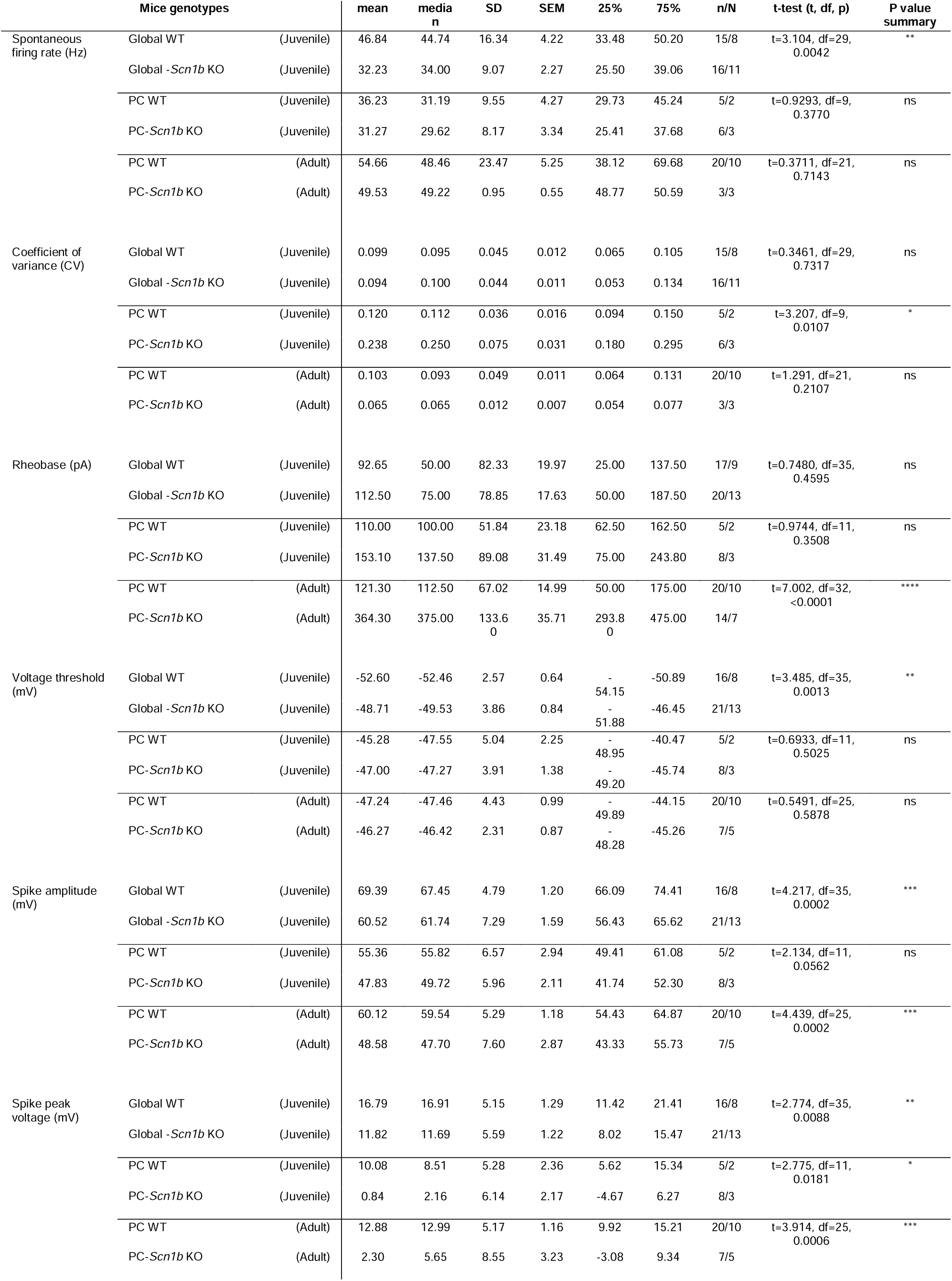
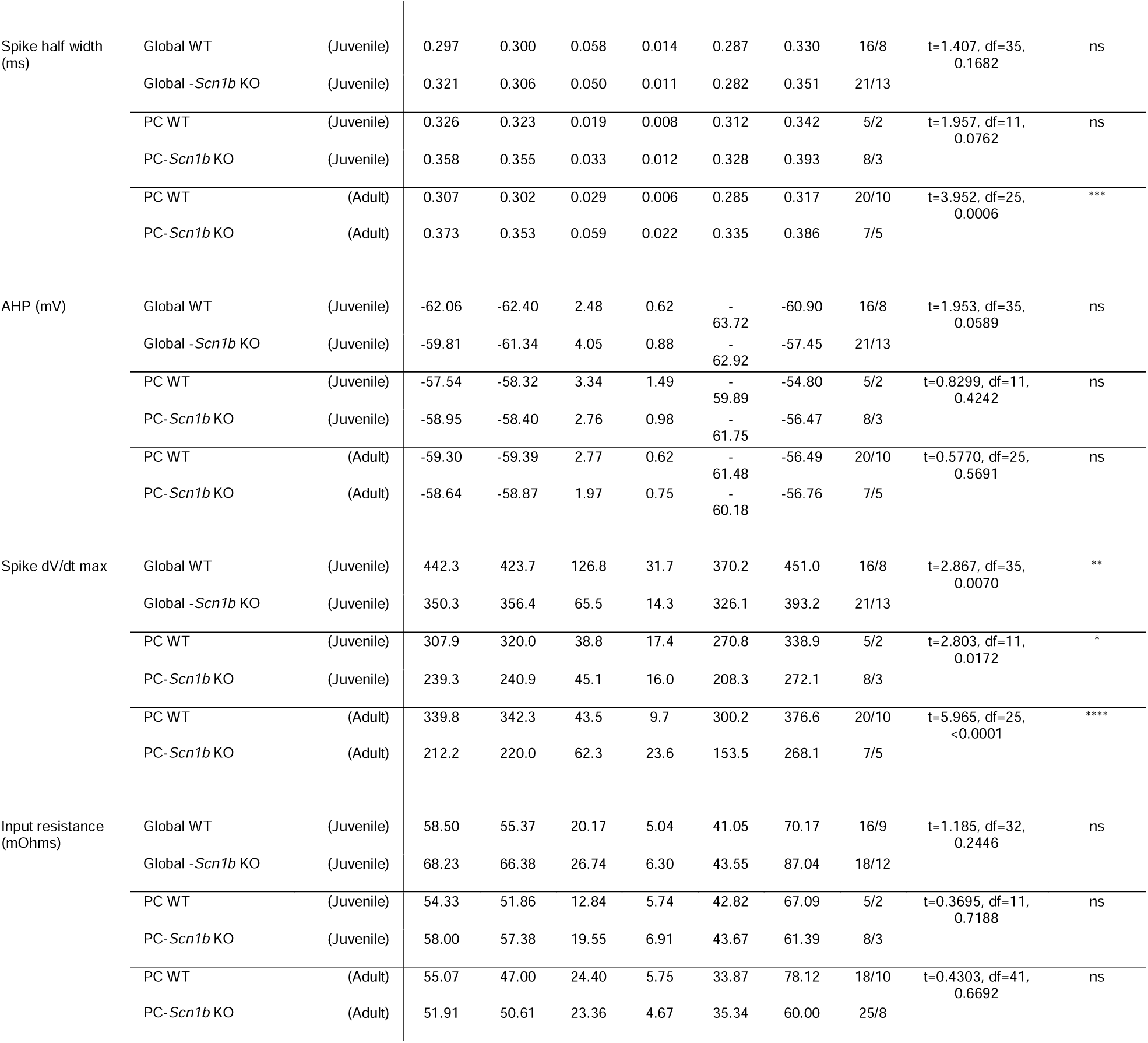
Descriptive statistics and t-tests. Columns show comparison made, experimental genotype, age, mean, median, standard deviation (SD), standard error of the mean (SEM), 25% confidence interval, 75% confidence interval, n/N (cell/mouse), t-test statistics, and p value. NS = not significant; *p<0.05, **p<0.01, ***p<0.0001, ****p<0.0001.

**Table 2:**
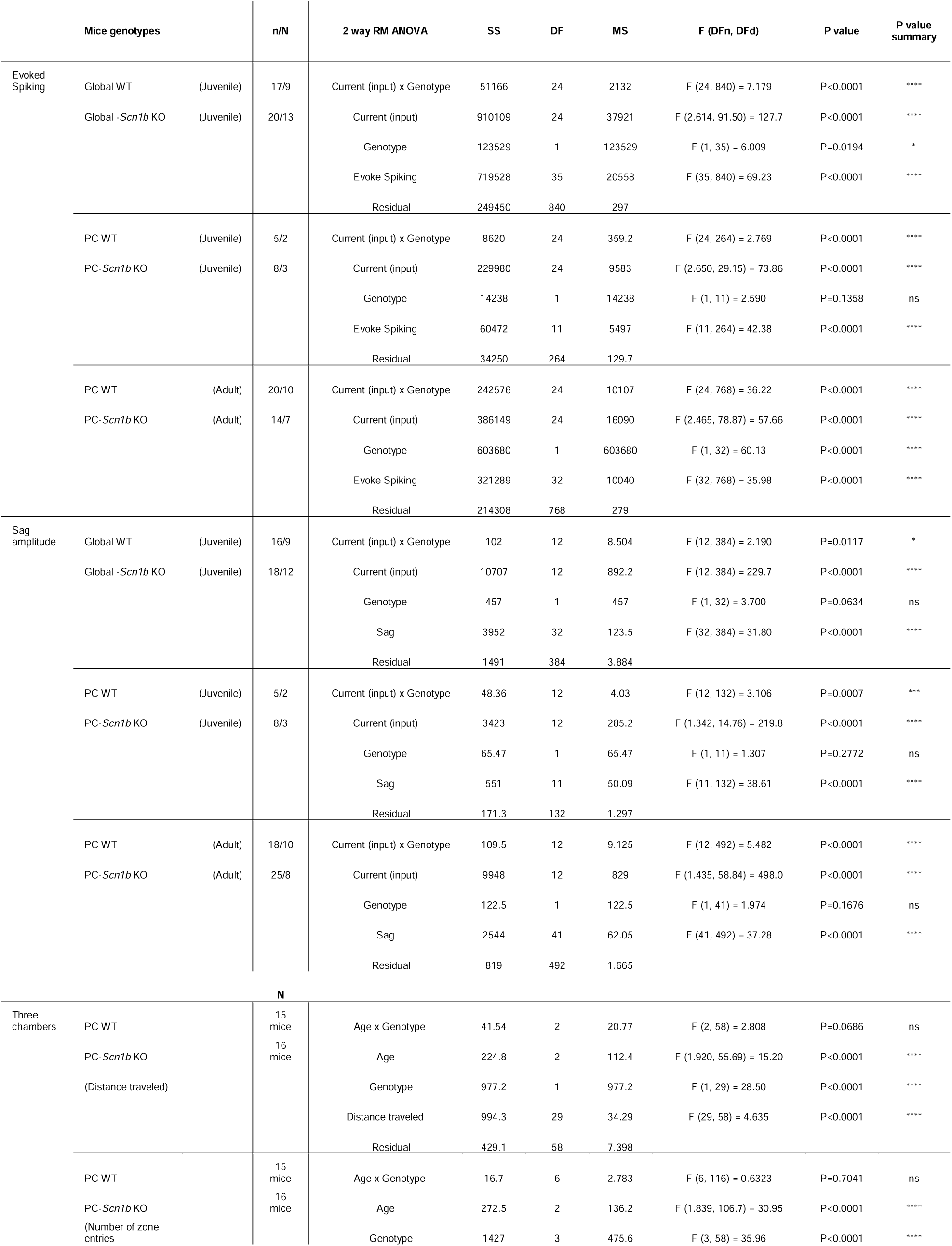
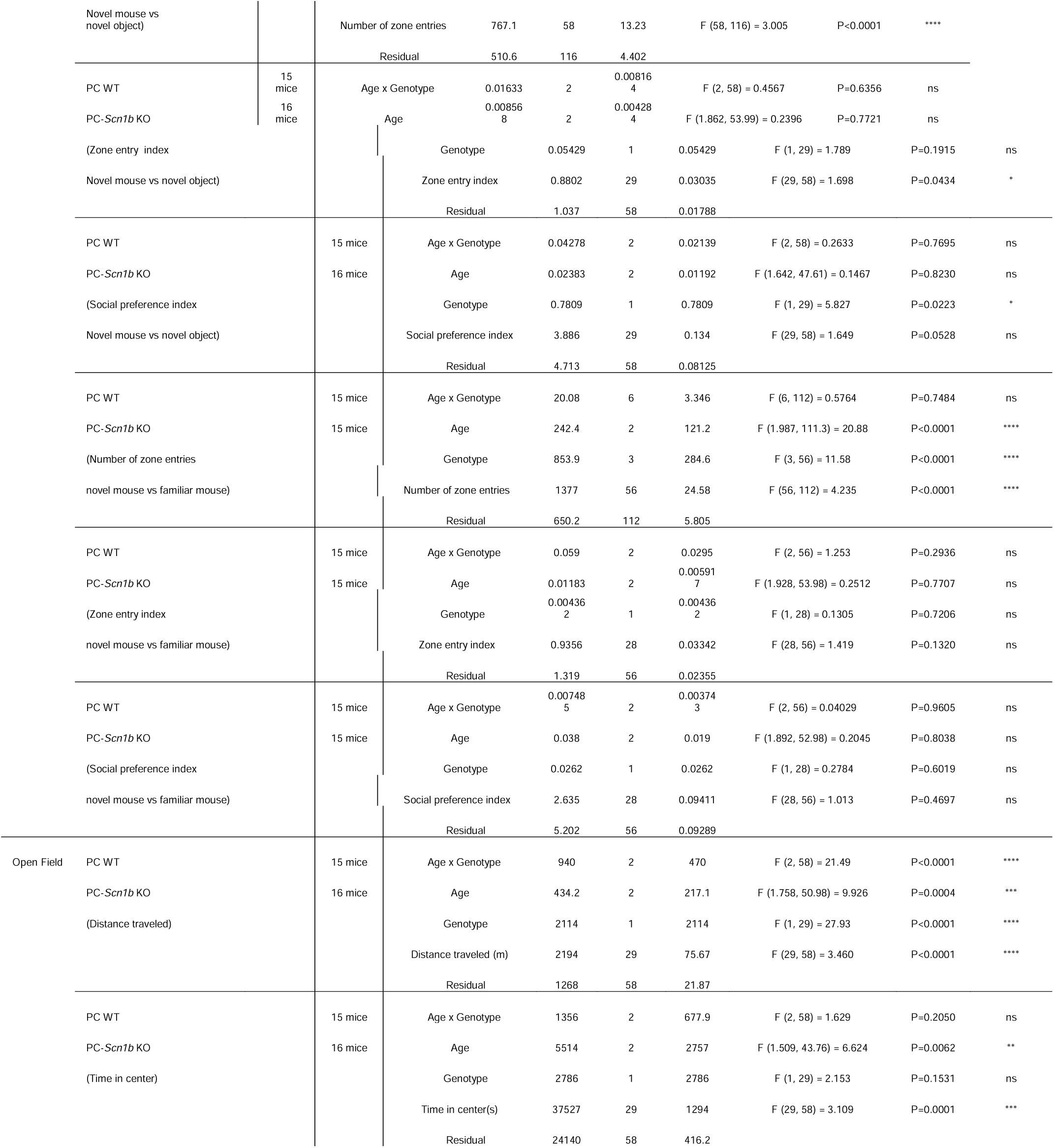
Descriptive statistics and ANOVAs. Columns show comparison made, experimental genotype, age, mean, n/N (cell/mouse), test type, sum of squares (SS), degrees of freedom (DF), standard deviation (SD), F values, p values. NS = not significant; *p<0.05, **p<0.01, ***p<0.0001, ****p<0.0001.

## Results

Results from juvenile Global-*Scn1b* KO, juvenile PC-*Scn1b* KO and adult PC-*Scn1b* KO mice were always compared to their respective WT littermates.

### Scn1b deficits cause changes to Purkinje cell spontaneous spiking

Cerebellar Purkinje cells continuously fire highly regular spontaneous action potentials at high rates (up to 100 Hz; Raman and Bean, 1999, Titley et al., 2020). This function allows downstream neurons to detect pauses in PC inputs, which is necessary for activation of plasticity mechanisms and cerebellar learning. Because *Scn1b* is involved in the specialized physiology of PCs (Aman et al., 2009) our first measure was spontaneous firing activity in PCs lacking *Scn1b*. We measured the spontaneous firing rate (Hz), and the coefficient of variation (CV) of the interspike-intervals (ISIs) to characterize the temporal structure of spike trains. The analysis was done over a 1 second duration of spontaneous firing recordings sampled at 10 kHz. Recordings were obtained across lobules of cerebellar vermis and paravermis slices (Fig. 1B, C). All WT PCs recorded exhibited spontaneous firing (with no injected current) (Fig. 1A). In the juvenile Global-*Scn1b* KO, all PCs exhibited spontaneous firing but mean spontaneous firing rate was significantly decreased (Fig. 1A, D, E; t-test; p=0.0042) with no change in the CV (Fig. 1F; t-test; p>0.05). In juvenile PC-*Scn1b* KO PCs, spontaneous firing was present in only 66% (6 out of 9) of neurons (Fig. 1D). In adult PC-*Scn1b* KO, the percentage of PCs exhibiting spontaneous firing decreased to 12% (3 out of 25). In PC- *Scn1b* KO brains, the PCs that did fire spontaneously maintained high firing rates statistically equivalent with WT (∼17 to 100 Hz; Fig. 1E; t-tests; p>0.05). The CV changed in the juvenile PC-*Scn1b* KO (Fig. 1F; t-test; p=0.0107) compared to their PC WT littermates, whereas no change was found in the adult PC-*Scn1b* KO vs adult PC-WT. To verify that PCs that did not exhibit spontaneous firing (or evoked firing as tested below) were indeed PCs and were otherwise high-quality recordings, we captured images of all cells and verified that resting membrane potential and input resistance were within normal ranges compared with WT recordings and previously reported values (Cupolillo et al., 2016). These findings suggest that *Scn1b* is necessary for maintaining the specialized physiology of PCs that supports a key aspect of their function.

**Figure 1:**
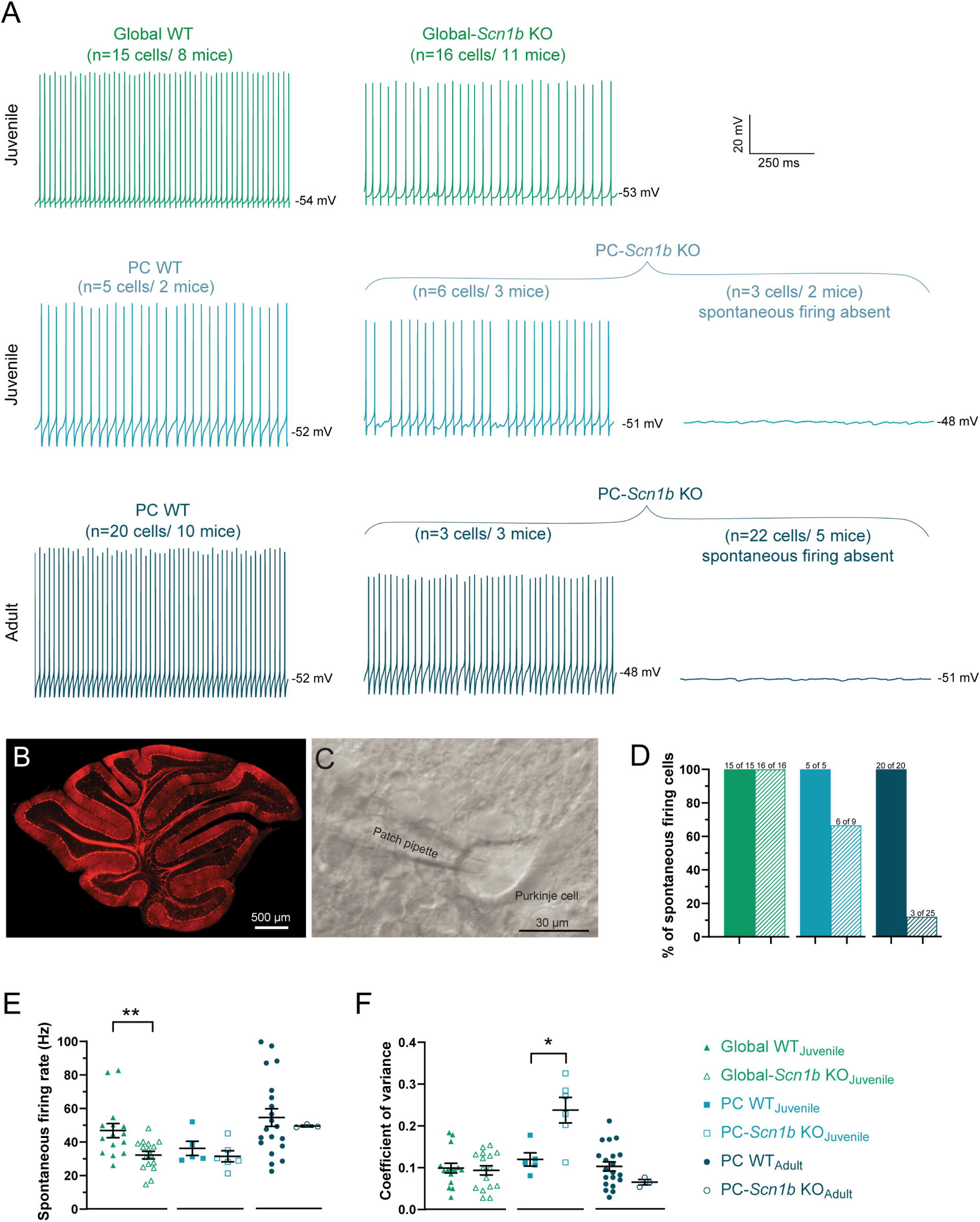
Spontaneous firing is decreased in *Scn1b*-deficient PCs. **A)** Representative spontaneous spike recordings from PCs from (middle panels) juvenile Global-*Scn1b* KO (green), juvenile PC-*Scn1b* KO (aqua blue), and adult PC-*Scn1b* KO (navy blue) and their respective wild-type littermates (left). Some cells from PC-*Scn1b* KO brains did not fire spontaneously (right). **B)** Cre-driven Tdtom fluorescence in an PC-*Scn1b* KO acute cerebellar slice (50 µm thick). **C)** Whole-cell current clamp recording of Purkinje cell soma in acute paravermis cerebellar slice (250µm thick). **D)** Percentage of PCs showing spontaneous firing cells by genotype. **E)** Spontaneous firing rate of PCs that exhibited spontaneous firing decreased in juvenile Global-*Scn1b* KO (open triangles; **p=0.0042; t-test) versus WT, but did not significantly change in PC-*Scn1b* KO neurons at either juvenile or adult. **F)** Coefficient of variation in firing rate is significantly increased in the juvenile PC-*Scn1b* KO (open squares; *p=0.0107, t-test) vs WT; juvenile Global-*Scn1b* KO (open triangles) and adult PC-*Scn1b* KO (open circles) were not different compared to WT littermates. Color scheme applies to all figures; n=cells/mice; *p<0.05; **p<0.01; solid black lines over scatter plots indicate mean±SEM for all figures.

### Evoked PC spiking is altered by loss of Scn1b

We next quantified the evoked firing rate of PCs in response to depolarizing current injections and measured the rheobase of the cell, which is the minimum injected current that results in an action potential. PCs were held at V_m_≅-65 mV using hyperpolarizing current injection to stop spontaneous activity, and 1-s duration depolarizing currents steps from 0 to 600 pA were injected in steps of 25 pA. Figure 2A illustrates representative evoked spiking recordings from *Scn1b*-deficient PCs (right) from juvenile Global-*Scn1b* KO (top), juvenile PC-*Scn1b* KO (middle) and adult PC-*Scn1b* KO (bottom), and their wild-type littermates (left). All Global-*Scn1b* KO PCs fired action potentials to depolarizing current injections. In juvenile PC-*Scn1b* KO 89% (8 out of 9) of the PCs exhibited evoked firing, while in adults only 58% (14 out of 24) of the PC-*Scn1b* KO PCs exhibited evoked firing (Fig. 2B). All WT PCs recorded exhibited evoked firing. Using only the neurons that did exhibit evoked spiking, we quantified spike input/output functions (evoked firing rate as a function of depolarizing current injection; Fig. 2C). Each *Scn1b*-deficient group exhibited a significant reduction in evoked action potentials compared to their wild-type littermates (two-way RM ANOVAs; p<0.0001). Rheobase increased significantly only in the adult PC-*Scn1b* KO group compared to PC-WT (Fig. 2D; t-test; p<0.0001). These results show that loss of *Scn1b* causes a progressive and severe loss of excitability in cerebellar PCs.

**Figure 2:**
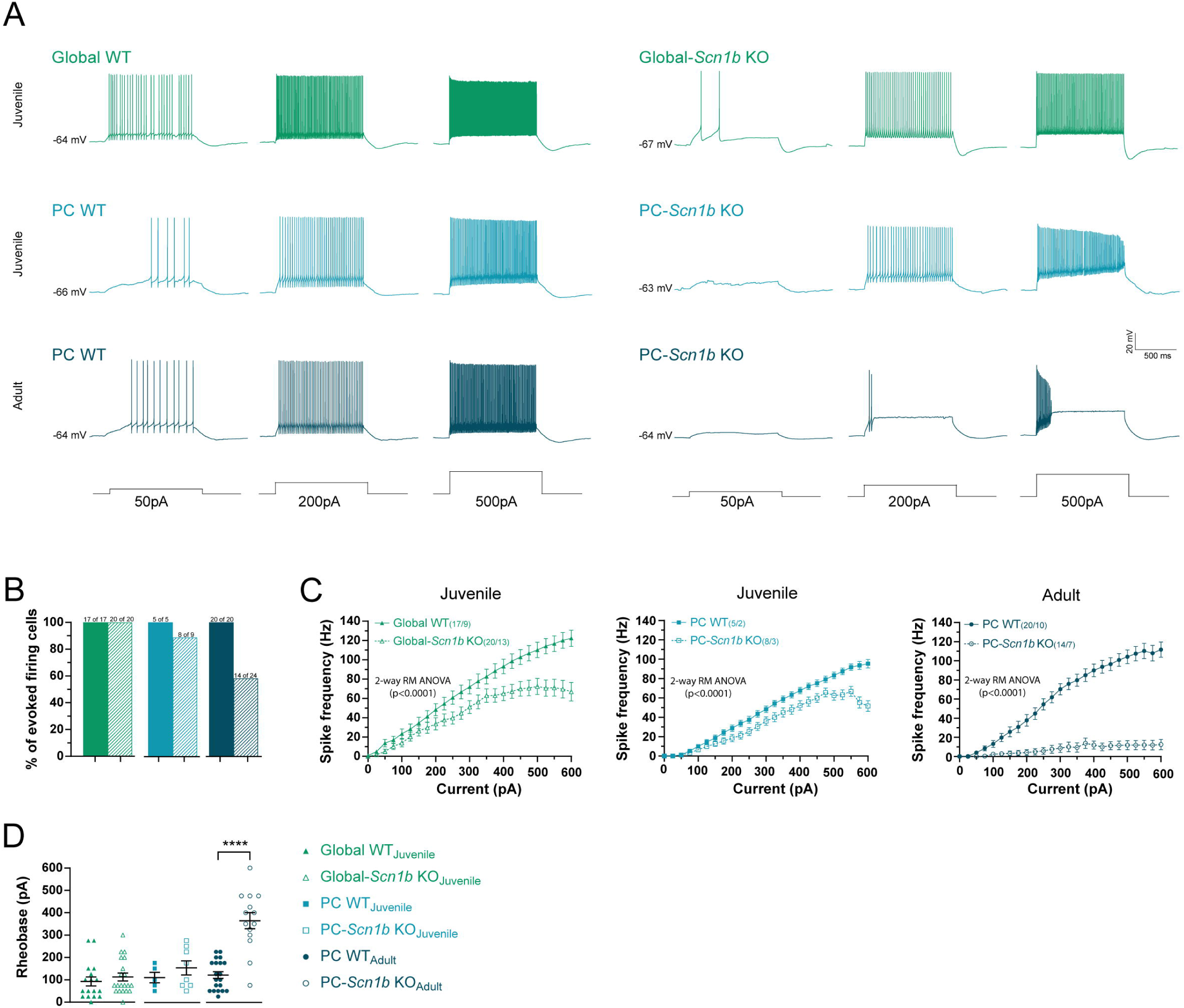
Evoked firing is decreased in *Scn1b*-deficient PCs. **A)** Representative evoked spiking recordings from *Scn1b*-deficient PCs (right) from juvenile Global-*Scn1b* KO (top), juvenile PC-*Scn1b* KO (middle), and adult PC-*Scn1b* KO (bottom), and their WT littermates (left). Cells were held at a hyperpolarized membrane potential, V_m_≅-65 mV, to stop spontaneous firing. Depolarizing current step amplitudes are shown at the bottom of each column of panels. **B)** Percentage of recorded cells exhibiting evoked firing cells by genotype. **C)** Evoked firing rate as a function of depolarizing current step amplitude for only neurons that exhibit spiking for each genotype. Two-way ANOVAs revealed significant decreases in firing in Global-*Scn1b* KO and PC-*Scn1b* KO fired fewer action potentials with increasing gradually depolarizing current injections (0 to 600pA) when compared to WT. **D)** Rheobase was significantly higher only in the adult PC-*Scn1b* KO compared to WT (t-test; ****p<0.0001).

### Loss of Scn1b alters spike and intrinsic properties in PCs

We studied the variability in the intrinsic electrical properties from singles spikes. All Global-*Scn1b* KO PCs evoked firing to a 300pA current injection, while in juvenile PC-*Scn1b* KO only 88% (8 out of 9) and in adult PC-*Scn1b* KO only 29% (7 out of 24) of the cells fired spikes to the stimulus (Fig. 3B). Using only the neurons that did exhibit evoked firing, we examined the spike intrinsic properties (peak voltage, threshold, spike half-width, AHP, spike amplitude and spike dV/dt max) in the first spike evoked at 300 pA injection. Results showed significant decreases in peak voltages in each *Scn1b*- deficient group when compared to their WT littermates (Fig. 3C). Voltage threshold depolarized in the Global-*Scn1b* KO (t-test, p<0.0013), but no changes were found in the juvenile or adult PC-*Scn1b* KO versus their WT littermates (Fig. 3D). Spike amplitude decreased in the Global-*Scn1b* KO (t-test, p<0.0002) and adult PC-*Scn1b* KO (t-test, p<0.0002), but no change was found in the juvenile PC-*Scn1b* KO when compared to their WT littermates (Fig. 3E). In the after-hyperpolarization voltage, we found no differences in the experimental groups when compared to the WT littermates (Fig. 3F). The spike half-width analysis showed no significant differences in the Global- *Scn1b* KO, or in the juvenile PC-*Scn1b* KO, but revealed an increase in the adult PC- *Scn1b* KO (t-test, p<0.0006) versus the WT (Fig. 3G). Finally, the maximum rate of change during the spike (max dV/dt) decreased in all groups lacking *Scn1b* (Fig. 3H). These analyses indicate that loss of *Scn1b* across models causes small action potentials with slower rise times (decreased max dV/dt) than in WT PCs. These results are highly suggestive of changes to voltage gated sodium channel function, in line with the eponymous function of *Scn1b* (Isom et al., 1992) and previous work specifically on *Scn1b* in PCs (Aman et al., 2009). That said, these data do not exclude the possibility that changes to the function of other ion channels may be causing or contributing, either directly through known *Scn1b*-ion channel interactions or indirectly through secondary mechanisms, to these physiological phenotypes.

**Figure 3:**
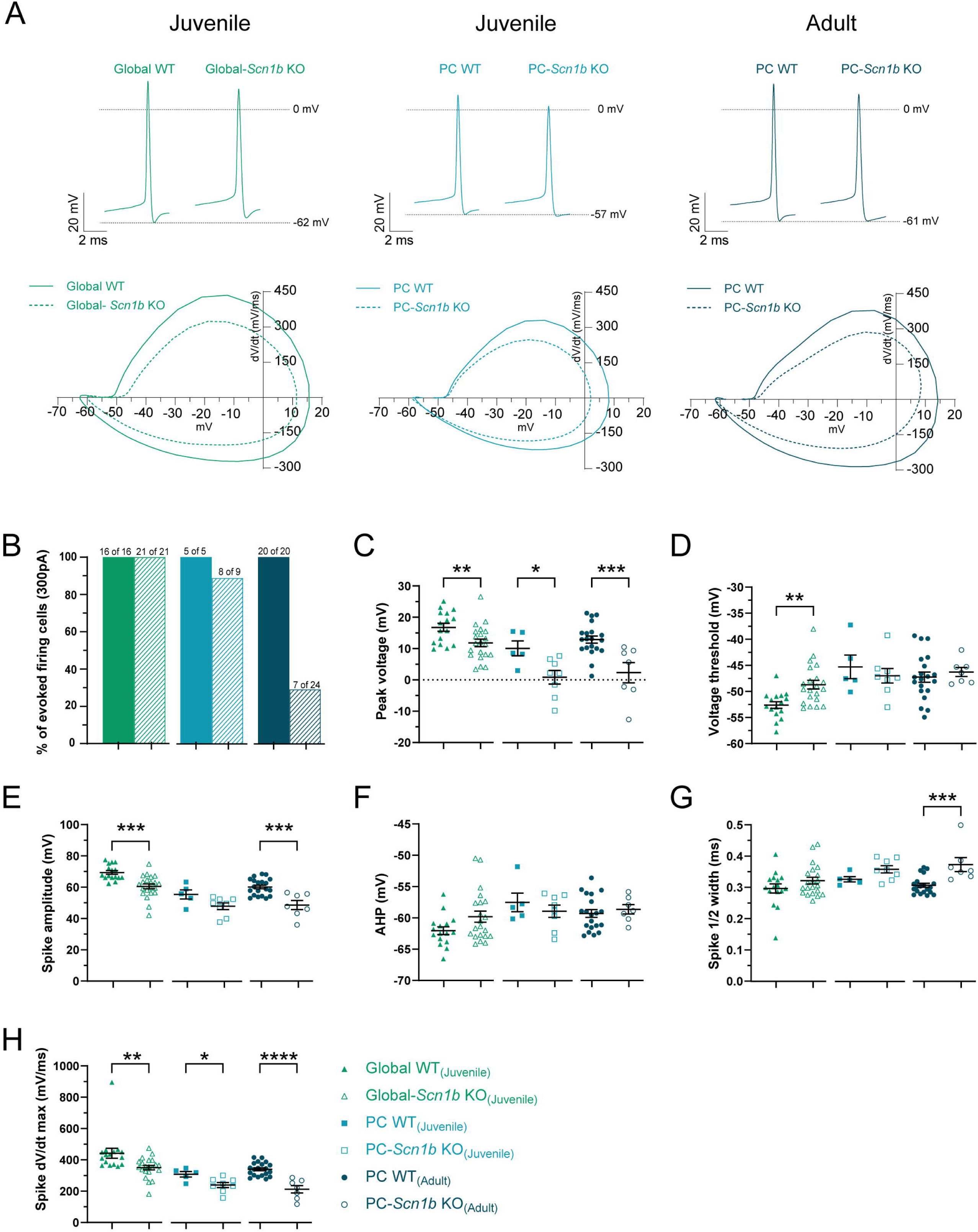
Spike intrinsic properties are altered in *Scn1b*-deficient PCs. **A)** For analysis of individual evoked spikes, we isolated the first spike fired in response to a 300-pA injection step. Top: Representative spikes from WT and experimental groups are illustrated side-by-side from juvenile Global-*Scn1b* KO (left), juvenile PC-*Scn1b* KO (middle), and adult PC-*Scn1b* KO (right). Dotted lines indicate zero mV and afterhyperpolarization peak. Bottom: phase plane plots are illustrated for the spikes above (WT: solid lines; *Scn1b*-deficient: dashed lines). **B)** Percentage of neurons recorded that fired spikes to this current step size, by genotype. **C)** Peak spike voltage was significantly reduced for each experimental group: juvenile Global-*Scn1b* KO (t-test; **p=0.008) and PC-*Scn1b* KO juvenile (t-test; *p=0.0181) and adult (t-test; ***p=0.0006, t-test) versus wild-type littermates. **D)** Voltage threshold significantly depolarized in the Global-*Scn1b* KO compared to WT(t-test; **p=0.0013). No significant differences were found between PC-*Scn1b* KOs at either age compared to WT littermates. **E)** Spike amplitude (peak voltage minus threshold) was reduced in Global-*Scn1b* KO (t-test; ***p=0.0002), not significantly different in the juvenile PC-*Scn1b* KO, and significantly reduced in adult PC-*Scn1b* KO (t-test; ***p=0.0002) compared with their WT littermates. **F)** Spike afterhyperpolarization voltage did not change significantly in any groups. **G)** Spike half width was significantly wider only in comparison between adult PC-*Scn1b* KO versus WT (t-test; ***p=0.0006). **H)** All three *Scn1b*-deficient groups showed significantly faster maximum spike dV/dt compared to WT (t-tests; Global-*Scn1b* KO: **p=0.0070; juvenile PC-*Scn1b* KO: *p=0.0172; adult Global-*Scn1b* KO: ****p=0.0001).

### Voltage sag increases in PCs lacking Scn1b

All Purkinje cells consistently showed a pronounced voltage sag in their response to hyperpolarizing current steps, which is indicative of hyperpolarization-activated cation current (I_h_). Sag was analyzed by calculating the amplitude of depolarization seen from the peak of a hyperpolarization response (600 pA current step) to steady state conditions (Fig. 4A). Input resistance (R_m_) was calculated from baseline voltage to the steady state voltage deflection in response to 50 pA hyperpolarizing current step. Voltage sag amplitude significantly increased in all groups lacking *Scn1b* compared with WT (Fig. 4B; RM-ANOVAs, p<0.05 for genotype X current interaction), while input resistance did not change when compared with their respective WT littermates (Fig. 4C; t-tests; p>0.05).

**Figure 4:**
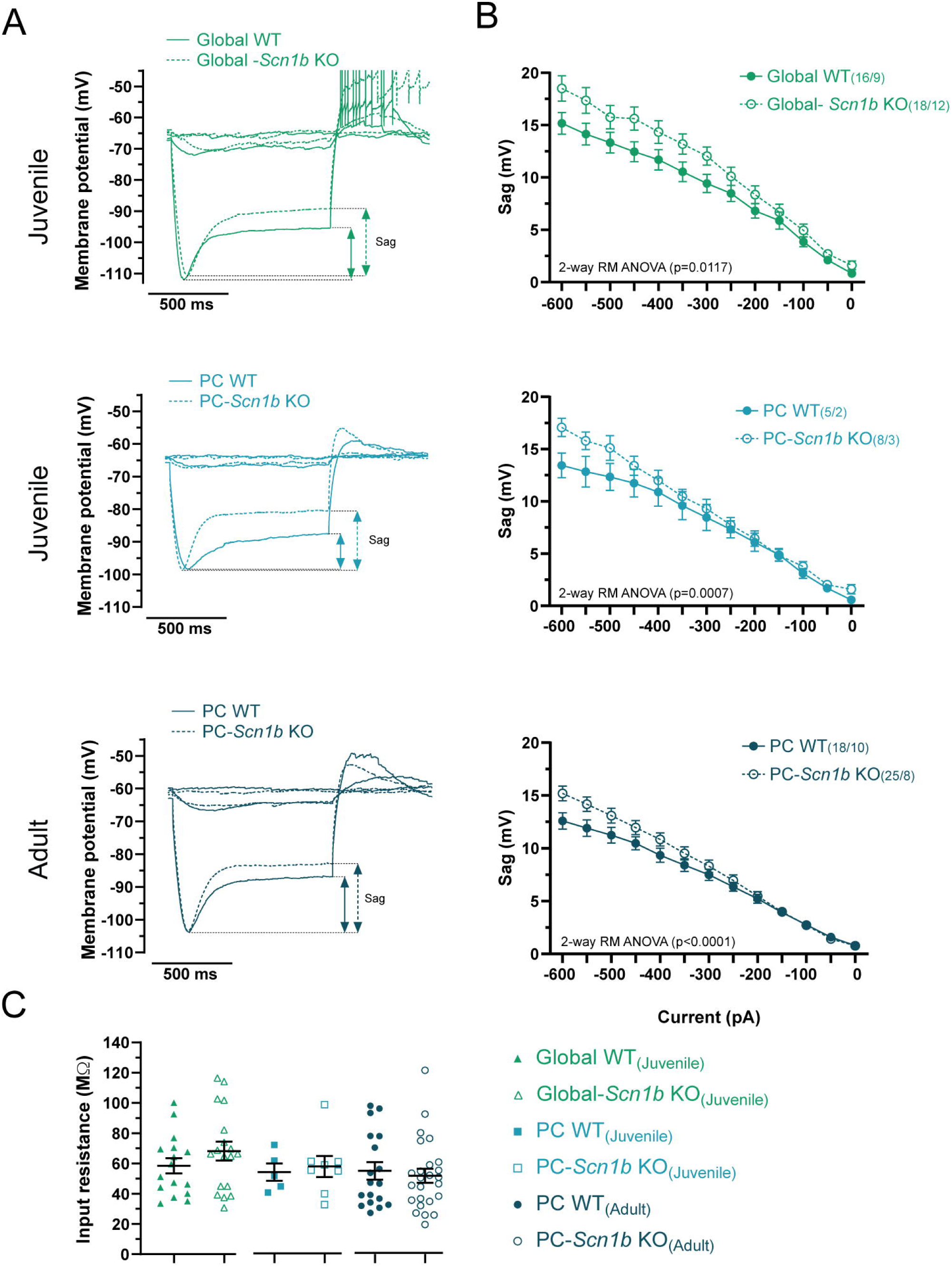
Voltage sag is increased but input resistance unchanged in *Scn1b*-deficient PCs. **A)** Voltage traces illustrating sag potentials (WT: solid lines; *Scn1b*-deficient: dashed lines) induced by hyperpolarizing current injections (-50pA and - 600pA) from juvenile Global-*Scn1b* KO, juvenile PC-*Scn1b* KO, adult PC-*Scn1b* KO, and their WT littermates. Vertical arrows represent voltage sag measurements. **B)** Voltage sag versus current injection plots showed an increase in amplitude in juvenile Global-*Scn1b* KO (open triangles; 2-way RM ANOVA, *p=0.0117), juvenile PC-*Scn1b* KO (open squares; 2-way RM ANOVA,***p=0.0007) and adult PC-*Scn1b* KO (open circles; 2-way RM ANOVA,****p<0.0001) compared with appropriate WT groups (inset p values are for voltage X genotype interactions from two-way repeated measures ANOVAs). **C)** Input resistances were not statistically different in cells lacking *Scn1b* when compared to WT littermates (t-test, p>0.05) (input resistance measure at -50pA current step). N=(cells/ mice).

### Cerebellar size and PC number are normal after loss of Scn1b

Some forms of cerebellar dysfunction involve loss of Purkinje cells and/or cerebellar degeneration. Additionally, major changes to neural activity levels can be a sign of changes in neuronal health and can induce cell death. Thus, we sought to ascertain whether loss of PCs was a part of the etiology of dysfunction following loss of *Scn1b*. We prepared cerebellar sections from adult (12 wk) PC*-Scn1b* KO and PC-WT mice for histology and confocal microscopy (Fig. 5). Sections were stained and imaged for Calbindin (Calb; green) and also imaged for expression of Cre-driven TdTomato (magenta). Using Calb staining, surface area of sagittal vermis sections was quantified (Fig. 5A-C). No significant difference was detected in cerebellar size between PC*-Scn1b* KO and PC-WT (t-test, p>0.05). PC density in the cell body layer was measured using higher magnification images of Calb staining (Fig. 5E). No difference in PC density found between genotypes (Fig. 5D; t-test; p>0.05). In PC*-Scn1b* KO sections, we compared Calb staining with Cre-driven TdTomato expression. Quantification of green/magenta colabeling (Fig. 5F) reveal a high degree of efficiency for both Calb labeling and Cre expression in PCs. Overall, unlike in some other forms of cerebellar dysfunction, PC cell death and cerebellar degeneration are not major contributors to altered phenotypes caused by loss of *Scn1b*.

**Figure 5:**
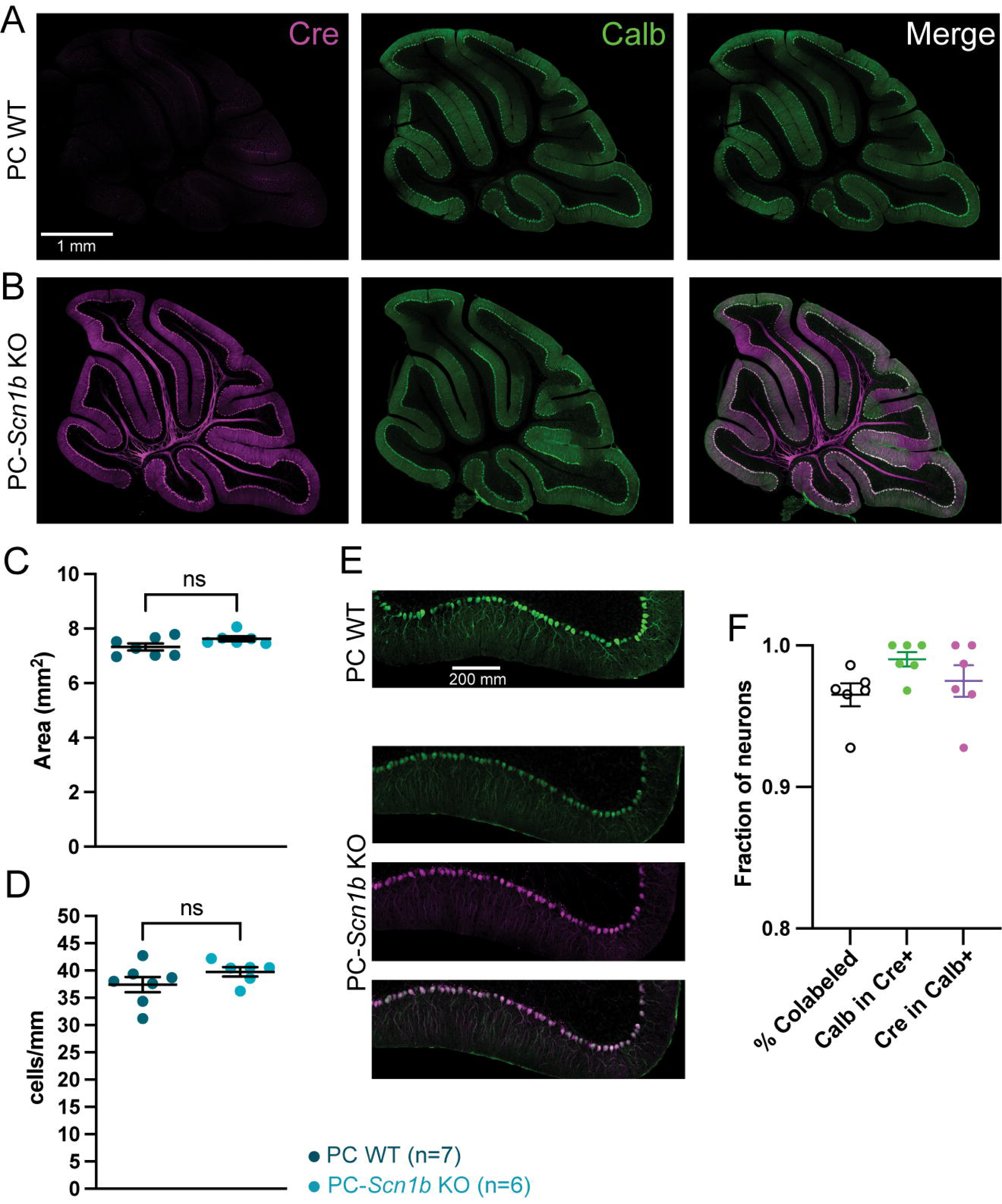
PC-*Scn1b* KO cerebellum shows no signs of degeneration. Confocal images showing Cre-dependent TdTom expression (left; magenta), IHC for Calb (green, middle), and merged images (right) for 12 wk WT (**A**) and PC-*Scn1b* KO (**B**) cerebellums. Scale bar in A applies to all panels of A and B. No significant differences were detected between groups for cerebellar area (**C**) or PC counts (**D**). Cell counts were made from higher magnification confocal images (**E**; scale bar in top panel applies to all panels in E). **F)** Colabeling was quantified in PC-*Scn1b* KO for Cre and Calb, Calb in Cre+ PCs, and Cre in Calb+ PCs.

### Scn1b loss causes decreased locomotor behavior

Movement disorders are one of the most common and negatively impactful manifestations of DEEs. *Scn1b* global knockout mice have visible ataxia, but ill health and early mortality make the extent and mechanisms of movement disorders due to loss of this gene difficult to study (Chen et al., 2004). Here we measured free exploration of an open field chamber to quantify locomotion in adult mice with PC-specific loss of *Scn1b*. The test was repeated at 8, 10, and 12 wk. Figure 6A shows representative tracks of mice exploring the open field. Over the five-minute test, PC*-Scn1b* KO traveled significantly less distance than their WT littermates (Fig. 6B; two-way RM ANOVA; effect of genotype: p<0.0001; effect of age: p=0.0004; genotype × age interaction: p<0.0001). WT mice exhibited a downward trend of distance traveled over repeated tests while PC*- Scn1b* KO mice did not, accounting for the significant effect of age and the genotype X age interaction. These quantifications illustrate the clearly observable deficit in motor behavior exhibited by mice with cerebellar *Scn1b* deficits.

**Figure 6:**
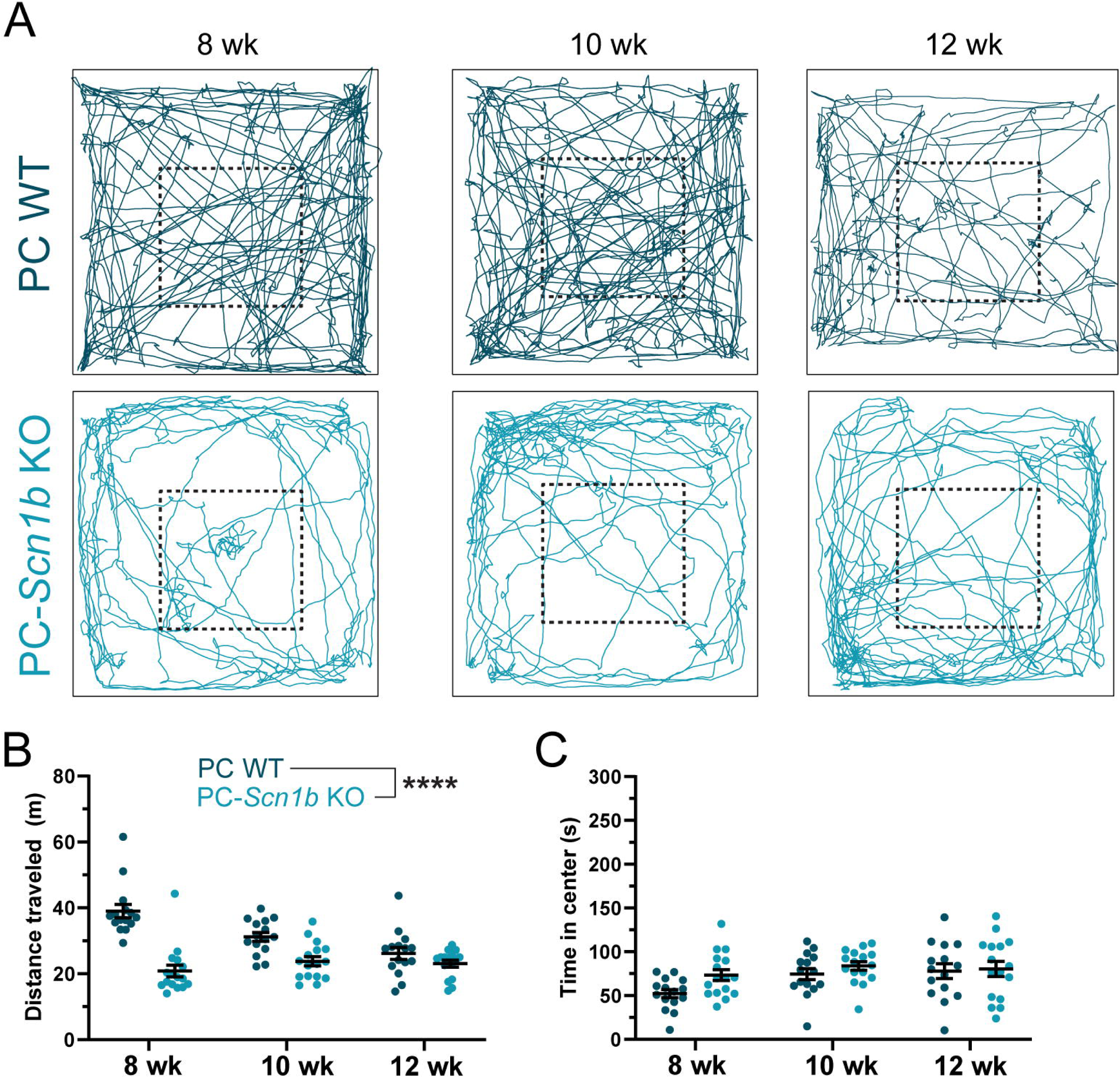
PC-*Scn1b* KO mice have locomotor deficits. **A)** Repeated motion tracking of a representative WT (upper) and a PC-*Scn1b* KO mouse (lower) at 8, 10, and 12 wk in a 50-cm^2^ open field arena over a 5-minute period. Dashed line illustrates 20-cm^2^ center zone. **B)** Total distance traveled was significantly less for PC-*Scn1b* KO mice compared with WT (two-way RM ANOVA, main effect of genotype). WT mice showed significant decreases in distance traveled across subsequent tests, PC-*Scn1b* KO mice did not (Tukey’s multiple comparisons tests). **C)** Time in center zone was not significantly different between genotypes (two-way RM ANOVA; ****p<0.0001, main effect of genotype, p>0.05).

From these tests we also quantified time spent in the central zone of the open field chamber. As prey species, mice tend to avoid open areas of unfamiliar environments compared with time spent near the walls of the chamber; time in center is a quantification of avoidance behavior. Analysis of these data revealed a significant effect of age, with both genotypes spending more time in the center zone over repeated tests, but no significant difference between genotypes or genotype × age interaction (Fig. 6C; two-way RM ANOVA; effect of genotype: p>0.05; effect of age: p=0.0062; genotype × age interaction: p>0.05). While the change in overall movement confounds these data to some extent, the lack of difference between genotypes suggests that PC-*Scn1b* KO mice are not more or less avoidant of the center of the open field than their WT littermates.

### Scn1b loss results in changes to social behavior

Autism Spectrum Disorder and/or autistic traits are among the most commonly reported neurological differences in Dravet syndrome and other DEEs. To further examine behavioral differences in PC-*Scn1b* KO mice, we used the 3-chamber test to examine social preference (Fig. 7A). Mice were placed in a divided arena with free access to an empty center zone (C), a chamber containing a caged novel mouse (M), and a chamber containing a caged novel object (O). Figure 7B illustrates representative tracking traces from repeated tests of a PC-*Scn1b* KO and a WT littermate at 8, 10, and 12 wk. Figure 7C shows all data of time spent in each chamber during these experiments. Analysis of social preference index ((time in M – time in O)/(time in M + time in O)), reveals a significant decrease in time spent with the novel mouse relative to novel object by PC- *Scn1b* KO mice compared with WT (Fig. 7D; two-way RM ANOVA; effect of genotype: p<0.0223; effect of age: p>0.05; genotype × age interaction: p>0.05). This indicates a decrease in social preference in PC-*Scn1b* KO mice. These data are complicated, however, by the difference in locomotion in PC-*Scn1b* KO mice described above, in that their ataxia may be a factor how much time they spend in each chamber. Figure 7E shows the total distance traveled by each mouse during the entire 5-minute three chamber test, again revealing a significant decrease in movement by PC-*Scn1b* KO mice compared to WT (two-way RM ANOVA; effect of genotype: p<0.0001; effect of age: p<0.0001; genotype × age interaction: p>0.05). We sought to understand if the decreased social preference in these mice was caused by changes to mobility in that mice were getting stuck in the chamber. Thus, we analyzed the number of entries mice made into the M and O chambers. PC-*Scn1b* KO mice showed fewer entries into both the M and O chambers compared to PC-WT (Fig. 7F; two-way RM ANOVA; effect of genotype p<0.0001; effect of age p<0.0001; genotype × age interaction p>0.05). However, despite differences in total entries, mice of both genotypes show an approximately equal number of entries into zone M as to zone O. This is reflected by the zone entry index [(M entries – O entries)/(M entries + O entries); Fig. 7G]. These data show that despite decreased mobility, PC-*Scn1b* KO chose to move in and out of the two chambers at equal rates, as WT mice do, but then spending less relative time in the chamber with the novel mouse. This suggests that the decreased social preference index is indeed due to a decreased social interest in mice with cerebellar *Scn1b* deficits.

**Figure 7:**
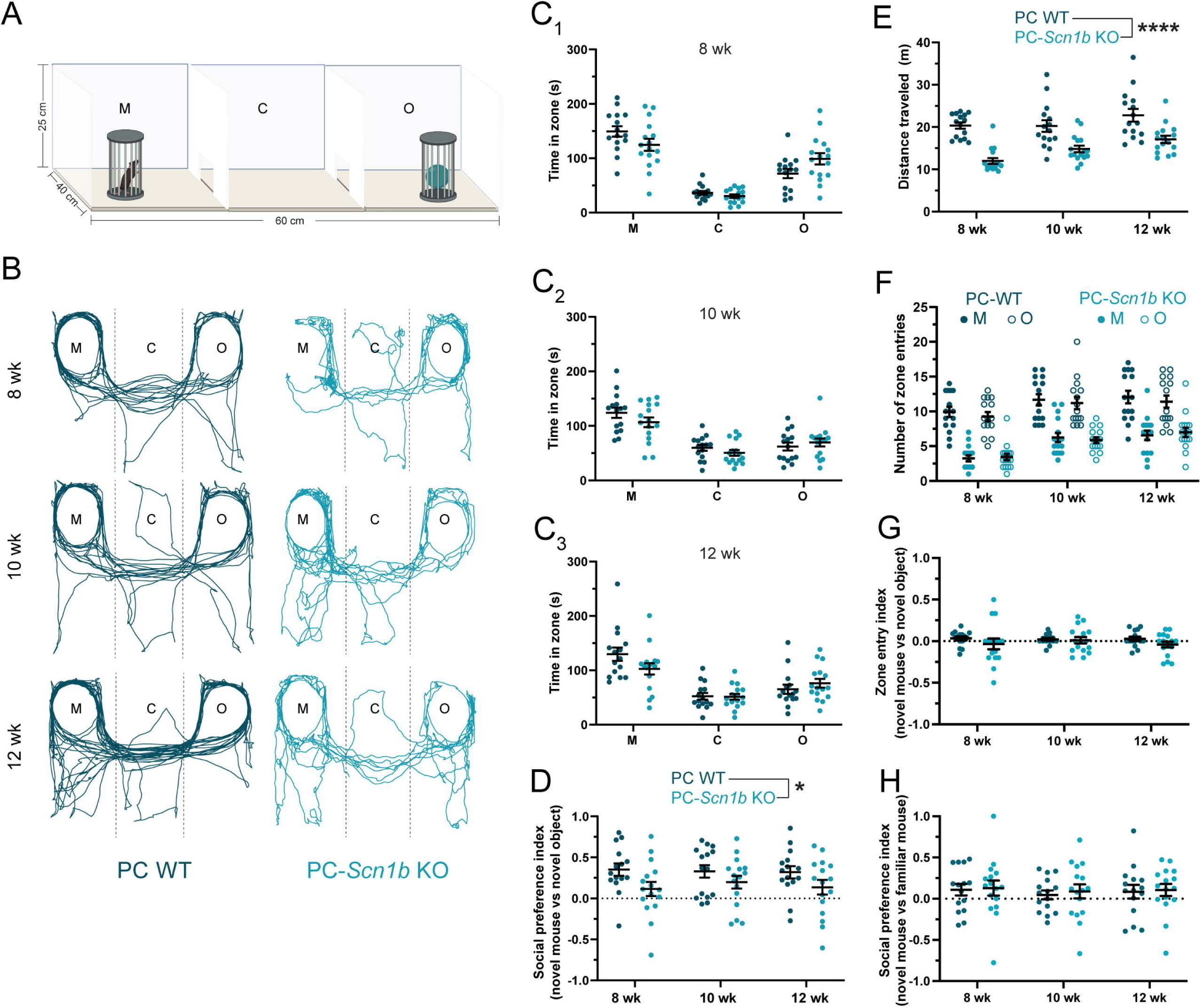
PC-*Scn1b* KO mice display altered social behavior. **A)** The three-chamber apparatus is a large box divided into mouse (M), center (C), and object (O) zones. The mouse and object are held under identical small cages. **B)** Representative trajectory plots of a PC-WT (left) and a PC-*Scn1b* KO (right) mouse, repeatedly tested at 8, 10 and 12 wk of age. **C)** Data for all mice showing time spent in each chamber over the 5-minute test period at each age. **D)** Analysis of social preference index revealed significant decrease in time spent in M relative to O zones in PC-*Scn1b* KO mice compared with WT (two-way RM ANOVA; effect of genotype: *p<0.0223). **E)** As in the open field, PC-*Scn1b* KO mice travelled significantly less total distance during the 3-chamber task than WT (two-way RM ANOVA; effect of genotype: ****p<0.0001). **F)** PC-*Scn1b* KO mice made fewer entries into both the M and O chambers during the 5-minute task than WT (two-way RM ANOVA; effect of genotype: ****p<0.0001). **G)** Index of entries into M and O zones was equivalent in both PC-*Scn1b* KO mice and WT mice. **H)** Phase three of the three chamber task revealed that both genotypes exhibited a mild preference (and statistically equivalent) preference for interaction with a novel mouse versus a familiar mouse.

Finally, we examined the effect of social novelty by testing preference for a familiar mouse in one chamber and a novel mouse in the opposite chamber (Fig. 7H). Both genotypes were equivalent in their social preference index for this test, showing slight increase in time spent with the novel mouse compared with the familiar mouse (two-way RM ANOVA; effect of genotype: p>0.05; effect of age: p>0.05; genotype × age interaction: p>0.05). Comparison of social preference index in this test revealed no significant differences between genotypes (Fig. 7H; two-way RM-ANOVA; p>0.05).

### PC-specific loss of Scn1b causes major deficits in eyeblink conditioning

Eyeblink conditioning is a Pavlovian associative learning paradigm that was first demonstrated in humans (Cason,1922) and later adapted for mice (Aiba, 1994; Chen et al., 1996). For the conditioned stimulus (CS) of this delayed eyeblink conditioning task we used a light flash that does not itself evoke an eyeblink. The CS was paired with an unconditioned stimulus (US), an air puff to the cornea which evokes eyelid closure. Over successive trials the pairing was learned, and a conditioned response (CR) was evoked: the mice began to close their eyelids in response to the CS, which peaked around the onset time of the US (Fig. 8A). This type of well-timed predictive eyeblink is a form of motor learning that is uniquely dependent on the cerebellum (McCormick and Thompson 1984; Garcia et al., 1999; Heiney et al., 2014). Here we found robust learning across trials in 5 of 6 PC-WT mice. This is quantified in two ways in each conditioning session: the proportion of trials (90 trials per session) in which animals showed a CR (CR rate) and the average fraction of eyelid closure (FEC) immediately before the US onset (Fig. 8B). When mice were tested with CS only, both these measures started at zero (i.e., no response) but increased and then plateaued over successive sessions (Fig. 8B, left). Conditioned responses were completely absent in all 5 PC-*Scn1b* KO mice (Fig. 8B, right). This finding indicates that loss of *Scn1b* in PCs disrupts normal plasticity/learning mechanisms in which the cerebellum is involved and suggests cerebellar dysfunction in the cognitive neurological disabilities of DEEs.

**Figure 8:**
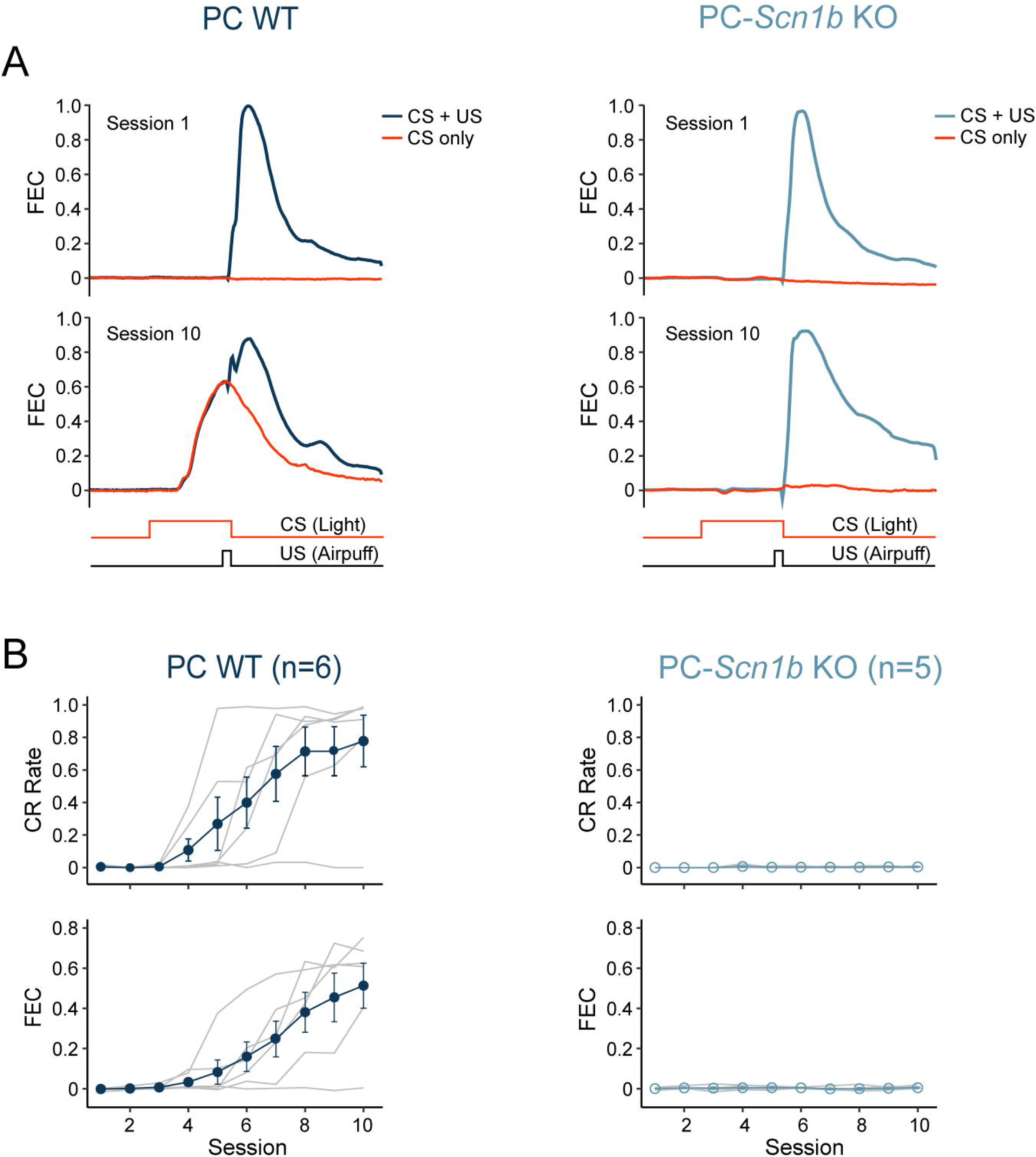
PC-*Scn1b* KO mice have cerebellar learning deficits. **A)** Delay eyeblink conditioning with a brief US co-terminated at the end of a longer CS (bottom traces). In session 1 (top panels), only US elicited eyeblinks (blue curves, average of CS+US trials). CS only trials did not elicit eyeblink (red curves, the average of CS-only trials). The curves indicate the fraction of eyelid closure (FEC) as a function of time during a trial. In session 10 (lower panels) WT mice exhibited a predictive eyeblink in response to CS-only trials; PC-*Scn1b* KO did not exhibit predictive eyeblinks. **B)** Mean CR rate (top) and FEC (bottom) show associative learning across sessions in the WT group (left) but not in PC-*Scn1b* KO (right). Gray lines represent individual mice.

## Discussion

*SCN1B* plays a diverse array of roles in brain development and adult maintenance of neural function. Loss of *SCN1B* causes severe developmental and epileptic encephalopathies (DEEs) that feature a broad range of neurological disabilities. Previous research on *Scn1b*, particularly using Global *Scn1b* KO mice, has provided insights into mechanisms by which gene products control neural development and cell physiology, but has been hampered by the severity of phenotypes and lethality of *Scn1b* loss, even with gene removal in adulthood and confined to specific neuron subpopulations. Our data validate a new model for examining *Scn1b* function and mechanisms underlying DEE-related neurological deficits. We found that loss of *Scn1b* restricted to cerebellar PCs did not lead to early mortality or systemic health issues, but caused decreased PC neuronal excitability, mirroring changes in the Global *Scn1b* KO, which progressively worsened into adulthood. We also found that removal of *Scn1b* in PCs was sufficient to induce significant deficits in motor, social, and cognitive processing. This work suggests mechanisms that may underlie some of the most negatively impactful neurological disabilities associated with *SCN1B*-DEE and a neurological localization upon which these disabilities converge (the cerebellum). Together, these suggest a potential target for developing treatments that could rescue functions in multiple neurological modalities.

The changes to excitability in PCs lacking *Scn1b* that we report are quite extreme, particularly in the adult mice. Both *in vivo* and *in vitro*, WT PCs fire ongoing trains of spontaneous action potentials, the frequency of which is modulated by synaptic input. PCs maintain this spiking behavior through coordinated action of specific ion channels. In particular, voltage gated Na_V_1.1 (*Scn1a*) and Na_V_1.6 (*Scn8a*) sodium channels are coexpressed with β1 and β4 (Raman et al., 1997; Raman and Bean, 1999; Aman et al., 2009). β subunits increase surface expression and alter gating and kinetics of sodium channel α subunits, but precise effects depend on cell type, subcellular compartment, and the combination of α and β subunits expressed (Bouza and Isom, 2017). In PCs, the specific α/β combination produces sodium currents with persistent and resurgent characteristics. Combined with expression of voltage-gated K^+^, Ca^2+^, Ca^2+^-activated K^+^, and HCN channels, these facilitate high frequency repetitive PC firing (Raman and Bean, 1999). Importantly, *Scn1b* gene products can directly interact with non-sodium channels important in regulating neural excitability (Deschênes and Tomaselli, 2002;

Marionneau et al., 2012; Nguyen et al., 2012). Further, while it is not known whether the phenotype is a primary phenotype caused by direct interactions with *Scn1b*, or secondary due to cell and circuit changes, increases in somatic HCN channel activity have been reported in neurons with *Scn1b* deficits, both in hippocampal CA1 pyramidal neurons (Chancey et al., 2023) and in PCs in the present study. Thus, while the single spike data presented here in PCs suggest changes to voltage gated sodium channel physiology, mixed mechanisms involving multiple ion channel families may be at play to produce the overall cell physiology phenotype. Further work will be required to delineate the ionic mechanisms underlying PC hypoexcitability after *Scn1b* loss.

Perturbations of the specialized aspects of PC physiology has been mechanistically linked to pathological outcomes. Classically, the cerebellum is best known for its role in motor coordination and balance. Variants in a diverse array of ion channel genes have been implicated in ataxia (for review, see Bushart and Shakkottai, 2019). Many genetic loss-of-function mouse models of ataxia genes exhibit altered locomotor behavior as well as altered PC electrophysiology. Some of these gene deficiency models exhibit PC hypoexcitability, others show cerebellar malformation or degeneration (Cendelin, 2014). Notably, ataxia can also result from changes to the regularity of spontaneous PC spiking. For example, variants in *Cacna1a*, encoding P/Q calcium channel α_1A_, and *Cacna2d2*, encoding the α2δ2 auxiliary subunit, can increase the coefficient of variance of repetitive PC firing without changing spontaneous or evoked spike frequency and result in substantial dyskinesia (Hoebeek et al., 2005; Walter et al., 2006). As in the human condition, global knockouts are likely to be causing physiological changes in other neurons in the cerebellar circuitry as well as beyond the cerebellum which could be major contributors to ataxic phenotypes. That said, PC-specific loss of ataxia-associated genes, such as *Cacna1a*, *Scn8a*, and *Klhl1*, is sufficient to cause ataxia (He et al., 2006; Levin et al., 2006; Todorov et al., 2012). Strikingly, PC-specific genetic replacement is sufficient to rescue both PC physiology and ataxia phenotypes in <u>global</u> *Kcnc3* knockout mice (Hurlock et al., 2008). In this context, our data showing changes to locomotor function after PC-specific loss of *Scn1b* make intuitive sense considering the PC hypoexcitability phenotypes we report.

The role of cerebellum in governing social behavior has been an area of growing research interest. Clinical findings have shown cerebellar abnormalities in postmortem studies of ASD (Fatemi et al., 2002) and high correlations between cerebellar anomalies and abnormal social behaviors (Moreno-Rius, 2019; Zhou et al., 2024). In mouse models, PC-specific knockout of genes linked to autism spectrum disorder is sufficient to cause decreased social interest in mice. This was first found in a *Tsc1*-deficient mouse model of Tuberous Sclerosis Complex (Tsai et al., 2012), in which a variety of autism-associated behaviors are exhibited after targeting gene deletion in PCs. Further analysis revealed that manipulating the activity of a small region of cerebellum, Right Crus 1, could induce changes to social behavior in wild type or rescue behavioral deficits in *Tsc1*-deficient mice (Stoodley et al., 2017). In another neurodevelopmental disorder mouse model, the *Fmr1* KO, targeting gene replacement to only the Crus 1 region was effective in improving behavioral measures in both the PC-specific and the global mutant model (Gibson et al., 2023). Physiological phenotypes in some, but not all, of these models also show a parallel to our work: PC hypoexcitability. PC-specific loss of *Tsc1* (Tsai et al., 2012), *Pten* (Cupolillo et al., 2016), and *Fmr1* (Gibson et al., 2023) all resulted in decreased spontaneous and/or evoked PC spike frequencies. It is notable that, unlike our study, none of these three models represents a channelopathy, i.e. pathology associated with a direct change to ion channel function. This suggests that decreases in PC output, due to any number of mechanisms such as direct or indirect changes in intrinsic excitability, altered expression of ion channels, or changes to synaptic function, may be a common mechanism linking gene variants, PC activity, and autistic features. This also raises the intriguing possibility that targeting increased PC excitability may be a generalizable therapeutic approach across multiple ASD etiologies.

The cerebellum also plays crucial roles in learning, memory, and cognition, including non-motor executive functions such as working memory (Lalonde et al., 1986) and cognitive flexibility (Pellegrino and Altman, 1979). Clinically, cerebellar lesions can result in cerebellar cognitive affective syndrome, characterized by a range of cognitive deficits (Schmahmann and Sherman, 1998). In animal models, eyeblink conditioning is a form of Pavlovian learning often used as a proxy for wider cognitive changes (Gormezano et al., 1962). The cerebellum is a vital node in the circuitry responsible for this form of learning (McCormick and Thompson, 1984; Garcia et al., 1999). Changes to eyeblink conditioning have been identified in a number of mouse models of genetic neurodevelopmental disorders (Kloth et al., 2015). We found that PC-specific removal of *Scn1b* completely abolished the expression of delay eyeblink conditioning in our mouse model. This parallels previous findings in DEE-associated models: PC-specific removal of *Scn8a*, which encodes a voltage gated sodium channel alpha subunit and is a DEE gene, disrupted delay eyeblink conditioning (Woodruff-Pak et al., 2006), altered PC sodium currents, and decreased PC spontaneous firing without inducing cerebellar malformations or cell loss (Levin et al., 2006). Other studies have shown that PC-specific removal of genes can disrupt eyeblink conditioning while having mixed effects on the physiology, morphology, and degeneration of PCs (Kim et al., 2011), including genes that can cause spinocerebellar ataxia (*Grm1*; Nakao et al., 2019), and autism spectrum disorder and learning disabilities (*Astn2*; Hanzel et al., 2024). Thus, the cerebellum provides a link between DEE genes and changes to learning, memory, and cognitive processing.

Through broad connections to other brain regions, cerebellar processing influences a diverse array of neural functions. PCs integrate inputs from cerebellar and non-cerebellar inputs and provide the sole output of cerebellar cortex information processing. PCs are both anatomically and physiologically specialized to fill this role. Our data show that the *Scn1b* gene, whose loss causes DEEs that include wide-ranging neurological disabilities, is a vital part of that physiological specialization. Both global and PC-specific deficiencies of *Scn1b* result in major changes to cellular function, motor, social, and cognitive behavior. Importantly, these data parallel previous reports of DEE genes that result in disorders combining disabilities of movement, social behavior, and cognition. DEEs are notorious for their medical intractability and negative impact on well-being; these results point to PCs as a potentially important target for development of treatments that could remedy a broad range of the neurological disabilities that impact the well-being of people with DEEs.

## Acknowledgements

We thank all the members of the Brumback-Howard labs for their help and insights throughout this project. We thank the Dravet Syndrome Foundation for inspiration for this project and ongoing support. We thank Lori Isom and lab for the gift of the *Scn1b* floxed mice. We also thank Jennifer Siegel, Richard Gray, and Raymond Chitwood for their help setting up the eyeblink conditioning system. Supported by funding from NIH/NINDS (NS112500), a Dravet Syndrome Foundation Research Grant, and a gift from the James and Katie Graham family.

